# Hi-C Contacts Encode Heterogeneity in Sub-diffusive Motion of *E. coli* Chromosomal Loci

**DOI:** 10.1101/2021.09.07.459285

**Authors:** Palash Bera, Abdul Wasim, Jagannath Mondal

**Affiliations:** Tata Institute of Fundamental Research, Hyderabad 500046, India

## Abstract

Underneath its apparently simple architecture, the circular chromosome of *E. coli* is known for displaying complex dynamics in its cytoplasm. Recent experiments have hinted at an inherently heterogeneous dynamics of chromosomal loci, the origin of which has largely been elusive. In this regard, here we investigate the loci dynamics of *E. coli* chromosome in a minimally growing condition at 30°C by integrating the experimentally derived Hi-C interaction matrix within a computer model. Our quantitative analysis demonstrates that, while the dynamics of the chromosome is sub-diffusive in a viscoelastic media in general, the diffusion constants and the diffusive exponents are strongly dependent on the spatial coordinates of chromosomal loci. In particular, the loci in Ter Macro-domain display slower mobility compared to the others. The result is found to be robust even in the presence of active noise. Interestingly, a series of control investigations reveal that the absence of Hi-C interactions in the model would have abolished the heterogeneity in loci diffusion, indicating that the observed coordinate-dependent chromosome dynamics is heavily dictated via Hi-C-guided longrange inter-loci communications. Overall, the study underscores the key role of Hi-C interactions in guiding the inter-loci encounter and in modulating the underlying heterogeneity of the loci diffusion.

## I. INTRODUCTION

The chromosome of the prototypical bacteria *E. coli* consists of 1.6 mm super-coiled circular DNA of 4.64 Mb. Unlike eukaryotes, it has no nucleus and it is confined within a (2 − 4)*µ*m long spherocylinder [1, 2]. However, the perception that bacterial chromosome is a randomly packed DNA, is fast changing. Precedent investigations have provided numerous evidences that the ‘nucleoid’ is just not a complex blob of genomic DNA, RNA and associated proteins. Rather it is being replaced by a picture of self-organized architecture with distinctly segregated macro-domains (MDs) and non structured (NS) regions [3, 4]. In addition to traditional molecular biology based experiments [3–9], the emergence of Chromosome conformation capture techniques in bacteria [10–12] are providing a microscopic view of the genome-level organization of its chromosome. In particular, a high resolution (5kb), Hi-C contact map for *E. coli* has recently been reported by Lioy et al [13]. The subsequent integration of this Hi-C interaction map in a polymer-based computational model [14], had bestowed a clear chromosomal compartmentalization, a fundamental building block of *E. coli* chromosomal architecture. In light of this, the center of our current investigation is: how does the self-organized nature of the *E. coli* chromosome impact the dynamics of its individual loci?

The dynamics of chromosomal loci is generally quantified by mean square displacements (MSD). Generally, for a diffusive particle *MSD* ∼ *τ*^*α*^, where *τ* is lag time and *α* is called MSD exponent. *α* = 1 indicates normal diffusion [15, 16], *α >* 1 indicates super-diffusion [17–20] and *α <* 1 implies sub-diffusion. Previous interesting experimental and theoretical investigations by Weber et al. [21–24] and Javer et al. [25] mainly focused on the spatiotemporal dynamics of *E. coli* chromosomal loci. Weber et al. explored the dynamics of chromosomal loci between (1 − 100)s time interval in LB medium (wt37LB). Their investigations showed that the movement of individual loci is sub-diffusive with a universal diffusive exponent *α* = 0.39 and this value was reported by them to be robust for both *E. coli* and *Caulobacter crescentus* with different drug treatments. A key conclusion of the investigations by Weber et al [21, 22] was that this universal value can not be explained by typical Rouse-like polymer model [26, 27]. Their measurement of velocity auto-correlation function with a negative peak indicated a viscoelastic nature of cytoplasm. A subsequent study by the same group had shown that ATP dependent biological activities do not change these robustness of MSD exponent but mobility of loci changes significantly [24]. However, in a later crucial investigation, Javer et al [25] had investigated the dynamics of a large set of chromosomal loci across the genome within very short time-scale ((0.1 − 10) second) in minimal medium, using high resolution tracking method. While this investigation also found that the MSD exponents of all loci indicate a sub-diffusive motion even in short time, there is a large variation of mobility across the chromosome loci. More importantly, the time scale and diffusivity of Ter macro domain were found to be very low compared to other macro domains. Together, these investigations bring out the complexity in the dynamics of the *E. coli*. chromosome and allude to a heterogeneous loci dependence. However, a microscopic picture of the origin of such observations are yet to emerge, especially at individual chromosome loci level.

Based on these precedent investigations and particularly the observation of the heterogeneous diffusivity across the loci, we surmised that the inter-loci cross-talk and specific identity of loci pair might play an important role. Towards this end, we planned on employing our recent model of *E. coli* chromosome [14], which had integrated the high-resolution (5 kb) Hi-C interaction map [13] in an excluded-volume polymer-based framework, for investigating the dynamics of the chromosomal loci. Specifically we use this Hi-C-encoded model to explore the mobility of each of 30 loci (same set of loci as investigated by Javer et al’s tracking experiments) via Brownian Dynamics (BD) simulations. Our analysis of the simulations trajectories reveal that both the diffusion constant and the exponents are loci-coordinate dependent and indicate a heterogeneous distribution. The result is found to be robust in presence of active noise. Interestingly, our control simulations via turning off the Hi-C interactions in the model abolishes this heterogeneity, suggesting that the loci-specific inter-genome interactions, as encoded in the Hi-C map, hold the key to the complex dynamics of the bacterial chromosome.

## II. RESULTS AND DISCUSSION

### A. An integrative Computer model Recapitulates E. coli’s Hi-C Contacts

The chromosome of *E. coli* grown in minimal media at 30°C (wt30MM) forms the basis of our investigation of the dynamics of its loci. Majority of the relevant experiments involving dynamics has also been performed on this growth condition. The cytoplasm of *E. coli* at this particular growth condition, contains a single chromosome, is not complicated by the fast replication process and hence is devoid of replication forks. As a result, the chromosome serves as a prototypical system which can be investigated for its short-time dynamics (20 min in real physical unit), without worrying about the chromosome segregation. Towards this end, we implemented a computer model of the *E. coli* chromosome by integrating beads-spring polymer topology with recently reported Hi-C interactions matrix of *E. coli* [13] chromosome at this particular condition (wt30MM). For this purpose, we employ a recently proposed protocol by our group [14]. As detailed in the *Simulation Model and Methods* section, the excluded volume interaction, Hi-C contacts and a spherocylindrical confinement form the key interactions of the model. We generated an ensemble of dynamical trajectories of the time-evolution of the chromosome model via BD simulations under over-damped condition (see *Model and Methods*). Figure 1 (a) renders a representative snapshot of the chromosome with color-coded encircled beads referring to various loci for different macro domains and various non-structured regions, as obtained in the simulations. The segregation of chromosome into six key macro domains, a signature feature of *E. coli*, is clearly evident. To assess the precision of the simulated model, we first compare the ensemble-averaged inter-bead contact probability with the experimental Hi-C matrix. Figure 1 (b) and 1(c) depict the heat maps of experimental and simulated contact probability matrices. The occurrence of an intense diagonal in both experimentally derived and simulated matrices represents that there is a higher contact probability between the chromosomal region for lower distances. Figure 1 (d) and 1 (e) demonstrate the heat map difference between experimental and simulated contact probability matrices and a histogram of this difference value respectively. A Pearson correlation coefficient value of 0.88 between experimental and simulated contact probability matrices and an absolute difference of the mean value of 0.067 between them imply reasonably good concurrence with in-vivo and in-silico chromosomal interactions.

**FIG. 1.**
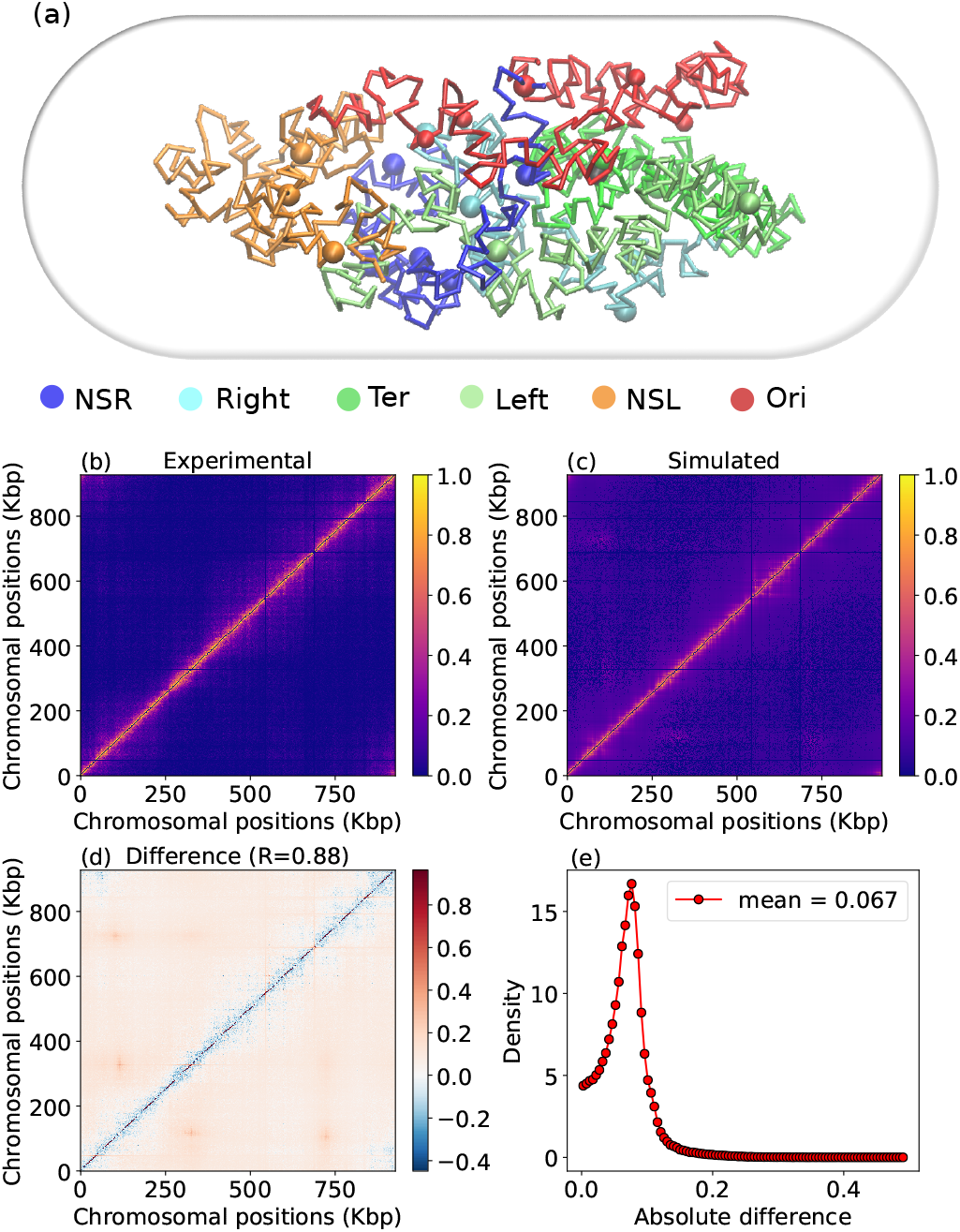
(a) Snapshots of an equilibrated chromosome, taken from a particular trajectory. The different color-coded regions represent corresponding macro-domains (MD) and non-structured (NS) regions and various colored coded encircled beads represent assorted loci for different MDs and NS regions. (b) Heat map of experimental Contact probability matrix. (c) Heat map of simulated contact probability matrix. This matrix is an ensemble average over 200 different initial configurations. (d) Heat map of difference matrix (simulated - experimental). R value is the Pearson correlation coefficient between simulated and experimental contact probability matrix values. (e) Histogram of absolute difference between simulated and experimental contact probability matrices with a mean of ∼ 0.067.

### B. Loci Mobility and Diffusive Exponents are Chromosome Coordinate dependent

Figure 2 (a) shows the genomic position of different loci in the circular chromosome [3, 4, 25]. Each of the color segments in Figure 2 (a) represents an individual macro domain and non structured region (NSR, NSL). Every macro domain and non structured region contains its own corresponding loci position, giving rise to a total number of 30 loci. To study the dynamics of chromosomal loci, here we have used BD simulations [28] in overdamped condition with a time step *δt* = 1*×*10^−4^*τ*_*BD*_ (see *Model and Methods* section). We simulate a time-range of (0.1 − 100)*τ*_*BD*_, which corresponds to (1.2*s* − 20*min*) (significantly shorter than the doubling time of the bacteria at 30°C in MM). First we have calculated the time averaged MSD for each trajectory (total 200 initial configurations) and subsequently have performed an average over all the trajectories, which we define as ‘ensemble average’ 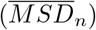. The time profiles of MSD and the MSD exponents enable us to characterize the type of diffusion i.e how fast the particles are spreading in space. The time-average MSD for a particular loci (n) defined as 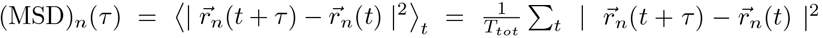, where *τ* and *T*_*tot*_ are lag time and total simulation time respectively. For a more appropriate power-law fitting of the ensemble average MSD, we have used 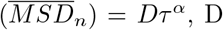 is constant). Generally 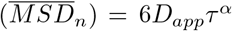 (in 3*d*), *D*_*app*_ is the apparent diffusion coefficient, so we denote *D* = 6*D*_*app*_ and *α* is the MSD exponent. We have divided the lag time into two regions: (0.1 − 10)*τ*_*BD*_ and (10 − 100)*τ*_*BD*_. We have also cross-checked that instead of the whole time lag, power law fitting is better in these two divided regions.

**FIG. 2.**
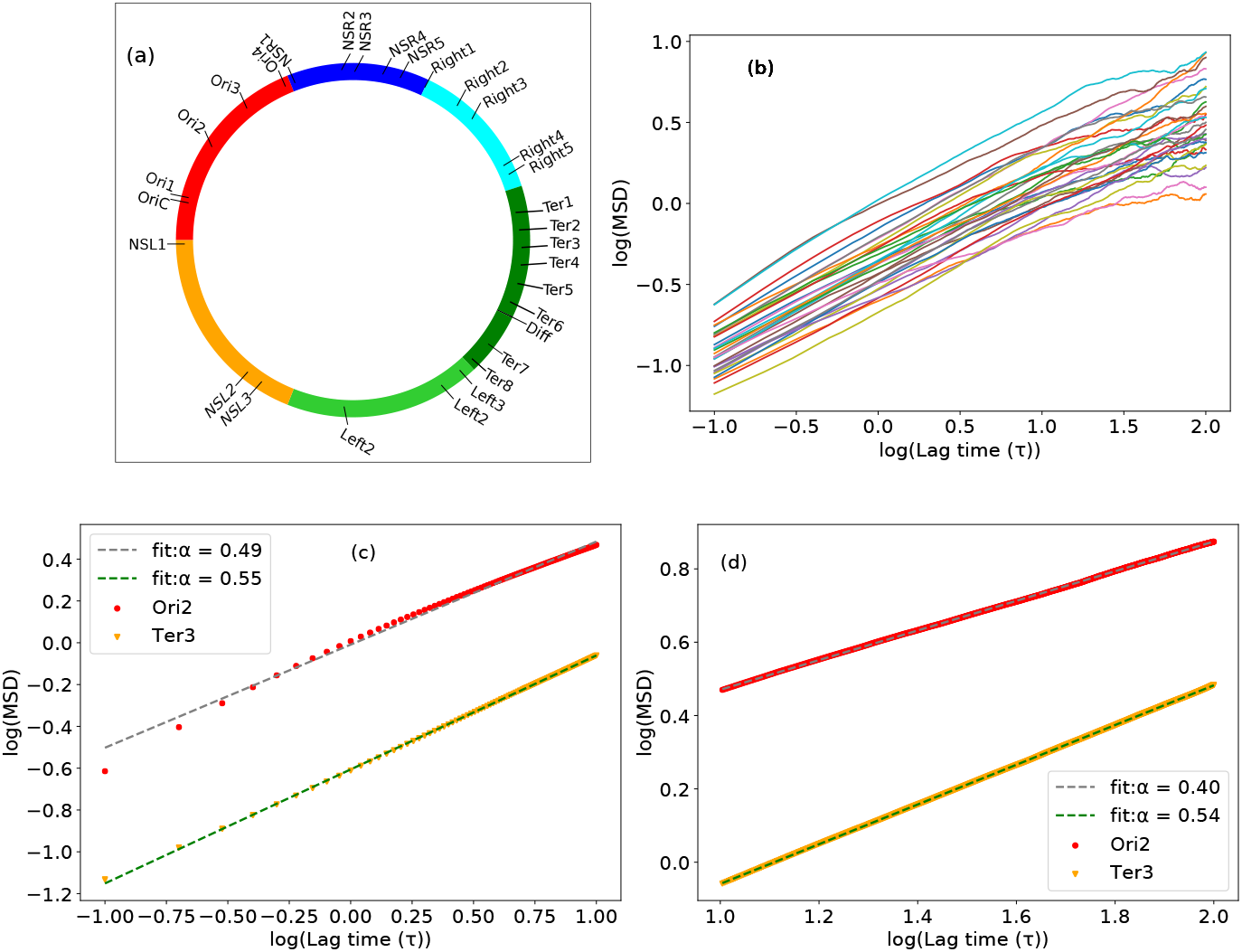
(a) Different loci position in a circular chromosome according to their genomic coordinates. Different color chunks represent the macro domains and non structured regions (NSR, NSL). (b) Time averaged MSD from single trajectory as a function of lag time for different loci. (c) Power law fitting of ensemble average MSD for two loci (Ori2 and Ter3) with time lag (0.1 − 10)*τ*_*BD*_. Scatter points (red for Ori2 and orange for Ter3) are simulated data and dotted lines (gray fit for Ori2 and green fit for Ter3) are fitted data. Ori2 and Ter3 have sub-diffusive exponents 0.49 and 0.55 respectively. (d) Power law fitting same as (c), with a lag time of (10 − 100)*τ*_*BD*_. Here Ori2 and Ter3 have MSD exponent 0.40 and 0.54 respectively, less than as mentioned in (c). For all the plots MSD and time are in units of *σ*^2^ and *τ*_*BD*_ respectively and these are in log scale.

Figure 2 (b) demonstrates the time averaged MSD, from a single trajectory as a function of lag time for different loci. The figure shows that there is a large spread in the time profiles of MSD across all loci. We first focus our attention on mobilities of two particular loci (Ori2, Ter3). Specifically, figure 2 (c) and 2 (d) compare the ensemble averaged MSD 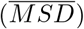 and its power law fitting for these two particular loci (Ori2, Ter3) at (0.1 − 10)*τ*_*BD*_ and (10 − 100)*τ*_*BD*_ respectively. In these two figures the scatter points are simulated data and the dotted lines are fitted data. From these figures, it is evident that there is a significant difference in individual mobility between Ori2 and Ter3 loci. To confirm this in-homogeneous mobility, we have also plotted the distribution of MSD values for a particular time 15*τ*_*BD*_ obtained from the time-averaged data of these two loci (figure S1(a) and S1(b) in SI). Together these figures depict that Ori2 and Ter3 differ notably in their displacement time profiles. More importantly, the individual power law fitting of the MSD profiles of these two loci reveal that the diffusive exponent is significantly lower than 1, thereby confirming their sub-diffusive behavior. More specifically, consistent with the loci-dependent trend in MSD, the power-law fitting indicates a significant difference in the exponent values between Ori2 and Ter3. For very short time lag ((0.1 − 10)*τ*_*BD*_) MSD exponents are given by: (*α* = 0.49 for Ori2 and *α* = 0.55 for Ter3, which are slightly higher compared to the long time lag ((10 − 100)*τ*_*BD*_) MSD exponents (*α* = 0.40 for Ori2 and *α* = 0.54 for Ter3). Experiment reports [25] that at shorter time scales, the lower and upper limit of these exponent as 0.4 and 0.5 respectively, which is in line to what we also observe from our simulations.

Together, the aforementioned observations of considerable difference in Ori2 and Ter3 movements reveal that both the loci mobility (characterized by MSD values) and the respective values of MSD exponents are significantly dependent on the relative coordinates of these two loci. These results prompted us for a comprehensive investigation of the mobility and the diffusive exponents for thirty chromosomal loci (see Figure 2 (a)) and characterize their trend. Accordingly, we calculated the MSD exponents and apparent diffusion constants from each time averaged simulation trajectories (total 30 *×* 200 = 6000) by fitting two different lag time regions ((0.1 − 10)*τ*_*BD*_ and (10 − 100)*τ*_*BD*_) and made a box plot and histogram plot for all of the loci. To get a more comprehensive insight into the loci-dependent behavior of mobility, figure 3 (a) and figure 3 (b) depict the box plot of apparent diffusion coefficient values of each of the thirty loci as a function of chromosomal coordinates, with lag time ((0.1 − 10)*τ*_*BD*_) and ((10 − 100)*τ*_*BD*_) respectively. From these two figures, it is clear that there are significant differences in diffusion coefficients that vary across chromosomal genomic positions in both time scales (short and long). In particular, it is also evident that among all the loci, Ori1 and Ori2 have substantially high diffusion coefficients and most of the Ter loci show lower diffusion coefficients. Figure 3 (c) and 3 (d) illustrate the box plot of diffusion exponents (*α*) of individual loci for short and long lag times respectively. For short lag time, the median values of the exponents lie higher than *α* = 0.5 and for long lag time this median values lies between *α* = 0.4 to *α* = 0.5 for most of the loci. This results are well corroborated with experimental findings by Javer et al [25], which reported that these median values lie around 0.4. In a similar fashion, figure 3 (e) and 3 (f) depict the histogram plot of MSD exponent for all trajectories of 30 loci for two different lag times. For short lag time, the mean value of the exponent *α*_*mean*_ ∼ 0.502 *±* 0.067 and for long lag time this value is lower *α*_*mean*_ ∼ 0.447 *±* 0.158. In both cases, the distribution of MSD exponent (*α*) values is very wide. Taken together, our Hi-C integrated computer model elucidates two key traits of *E. coli* chromosome : a) slow sub-diffusive dynamics and b) heterogeneous chromosomal loci mobility. These results are consistent with previous experiments, [21, 25] and prompt us to elucidate the underlying mechanism of these features.

**FIG. 3.**
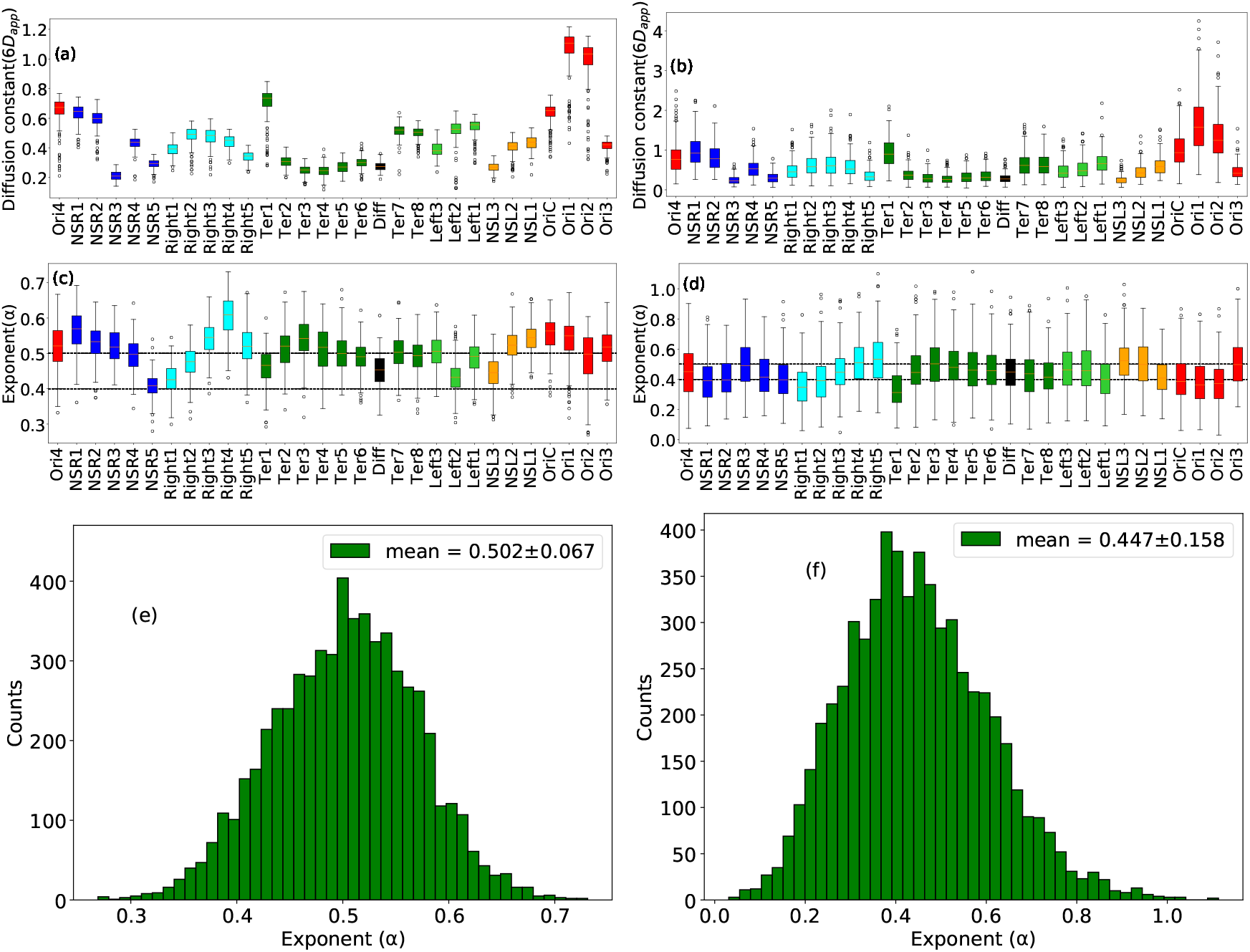
(Box plot of Diffusion constant as a function chromosomal loci in presence of Hi-C with lag time (a) (0.1 − 10)*τ*_*BD*_ and (b) (10 − 100)*τ*_*BD*_. In both case the value of diffusion constants vary from loci to loci, so mobility of chromosomal loci depend on their genomic co-ordinate. Box plot of MSD exponent for each and individual loci, by taking MSD exponent from individual trajectory of loci, with lag time (c) (0.1 − 10)*τ*_*BD*_ and (d) (10 − 100)*τ*_*BD*_. For most of the loci median values of the exponent lies higher than *α* = 0.5 for short time lag and between *α* = 0.4 to *α* = 0.5 for long time lag respectively. Histogram of MSD exponent by taking all of the trajectory of 30 loci (total (200 *×* 30 = 6000 trajectory) with lag time (e) (0.1 − 10)*τ*_*BD*_ and (f) (10 − 100)*τ*_*BD*_. From the histogram mean vale of the exponent is *α*_*mean*_ ∼ 0.502 *±* 0.067 for short time lag and *α*_*mean*_ ∼ 0.447 *±* 0.158 for long time lag respectively. Taken together four figure we can say chromosomal dynamics are heterogeneous with respect to individual loci as well as time.

### C. Hi-C contacts as the origin of the heterogeneity in *E. coli*. chromosomal diffusivity

The preceding discussions pointed out to a significant loci-coordinate-dependent sub-diffusive motion of chromosome, largely consistent with the previous measurements by Javer et al [25]. These experimentally consistent observations of heterogeneous mobility across chromosomal loci in our computer model demands a more microscopic physical interpretation. We noted that one of the key features of the current model, that sets it apart from other excluded-volume interaction model or other generic polymer-based ‘Rouse model’, is the integral presence of ‘Hi-C’ interaction potentials to capture the experimentally obtained inter-beads contact matrices. We speculated that these Hi-C contact might have a role in dictating the loci-dependent heterogeneity. Accordingly, to dissect its specific role, if any, we performed a fresh set of control dynamical simulations by removing Hi-C contacts from our computer model.

Figure 4 (a) and 4 (b) plot the MSD exponents of all loci, as derived from the simulations trajectories of the model in absence of Hi-C interaction potential, at short and long two lag times. For the purpose of comparison, we also reproduced the MSD exponent of same set of loci as derived in simulations with Hi-C contacts (figure 4 (c) and 4 (d)). The comparison suggests that the MSD exponents are uniformly similar across all loci in absence of Hi-C contacts. However, turning on these interactions in the model recapitulates the loci-dependence of MSD exponents. On a similar spirit, to investigate this loci-specific heterogeneity in diffusion coefficients, we have also calculated this diffusion constant value from simulated trajectories in the absence of Hi-C interactions. Figure S2(a) and Figure S2(b) demonstrate the apparent diffusion coefficient values as a function of chromosomal coordinates, in the absence of Hi-C interactions with lag time ((0.1 −10)*τ*_*BD*_) and ((10 −100)*τ*_*BD*_) respectively. When compared with the results of simulated data in presence of Hi-C contacts (Figure S2(c) and Figure S2(d)), it is obvious that without Hi-C contacts, all the loci display nearly the same diffusion constant values, with two different time regions. In combination, these analysis, based on control simulations, dissect the decisive role of specific Hi-C contacts in capturing the inherent loci-dependent mobility.

**FIG. 4.**
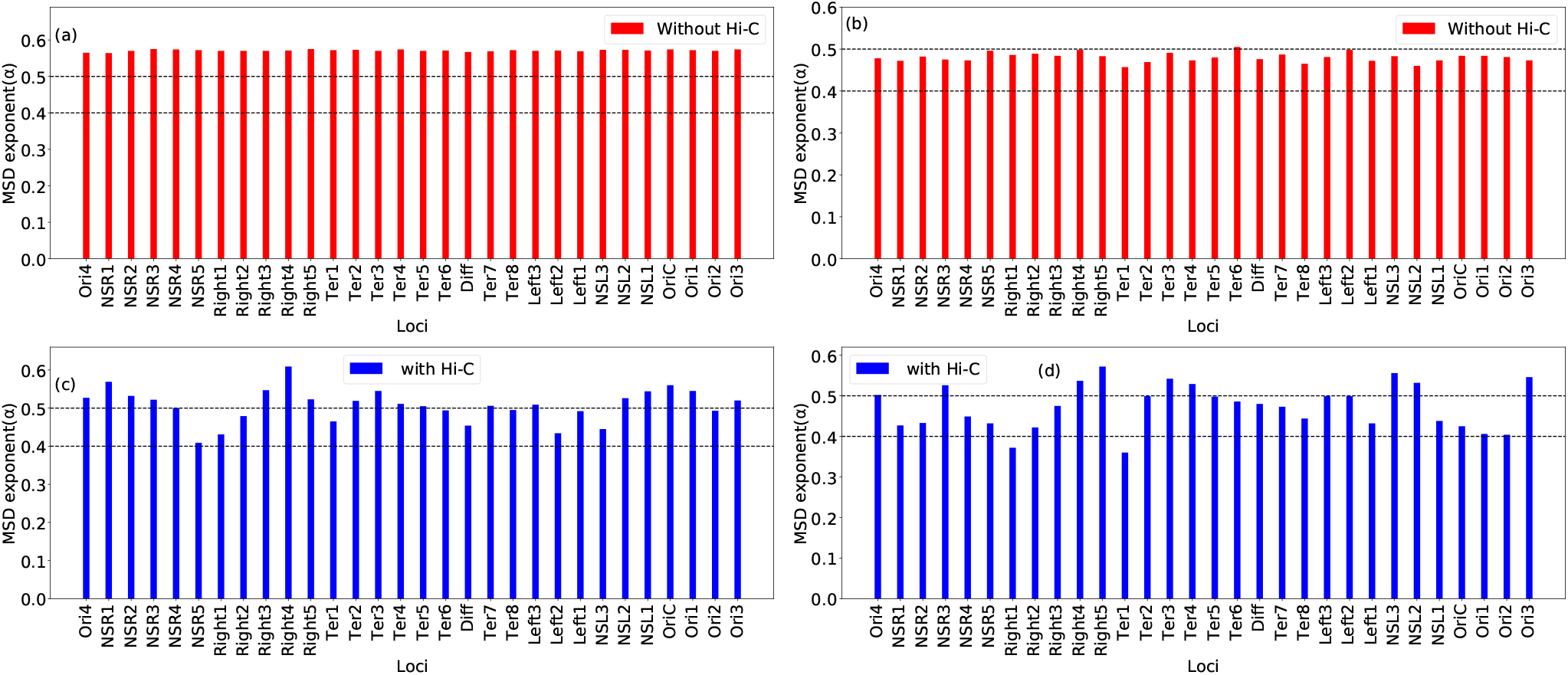
Bar plot of MSD exponent as a function of chromosomal loci in absence of Hi-C with lag time (a) (0.1 − 10)*τ*_*BD*_ and (b) (10 − 100)*τ*_*BD*_. MSD exponents are almost same with respect to different loci for both cases. Bar plot of sub-diffusive exponent as a function chromosomal loci in presence of Hi-C with lag time (c) (0.1 − 10)*τ*_*BD*_ and (d) (10 − 100)*τ*_*BD*_. There is a clear heterogeneity in sub-diffusive exponents with respect to different loci. The exponents are slightly higher for small time lag as compared with higher time lag.

The resolution of our model and the explicit incorporation of the Hi-C interaction potential allow us to track specific inter-loci contacts and pinpoint the underlying reason for the coordinate-dependent loci dynamics. In the current protocol, the inter-genomic distance is modeled as inversely proportional the Hi-C contact probability (*D*_*ij*_ = *σ/P*_*ij*_). Accordingly, the number of inter-beads contacts connected to each loci gets modulated. In particular, Ori1 and Ori2 loci show one of the higher mobilities, while most of the Ter loci (Ter2, Ter3, Ter4, Ter5, Ter6), Diff, NSR3 and NSL3 show the slowest mobility. On the other hand, some loci (NSR3, Right4, Right5, Ter3, Ter4, NSL2, NSL3 and Ori3) are showing higher exponent (*α >* 0.5) values and some of them (Right1 and Ter1) are showing lower exponent (*α <* 0.4) value, but the values of these exponents lie around *α* ∼ 0.44 *±* 0.158, for longer lag time.

To get a closer insight on why Ter macro-domain (MD) demonstrates comparatively lower mobility compared to other MDs, we have computed the fraction of connections between different MDs and NS regions in our model. Figure S3 highlights the heat map of the fraction of Hi-C contact formed within MDs and NS regions. This suggests that the fraction of intra-chromosomal connections within the Ter MD is relatively higher compared to other MDs, which slows down the mobility of Ter and its loci. On a related note, prior investigations [6, 29] refer that the presence of nucleotide associated protein MatP in the Ter domain plays an important role in insulating it from the other MDs and making it more compact [30]. We wanted to explore if the current model can predict the role that MatP would have played in modulating the loci dynamics of *E. coli*. Accordingly, we made use of Lioy et al’s experimentally measured Hi-C data of mutated bacteria devoid of MatP [13] and performed a fresh set of simulations for capturing the dynamics of MatP-devoid chromosome (ΔMatP). (see *SI Model and Methods*) Figure S4(a) and S4(b) depict the diffusion constant values of all loci with wild type (WT) and ΔMatP mutant at two different lag times, (0.1 − 10)*τ*_*BD*_ and (10 − 100)*τ*_*Bd*_ respectively. For mutant MatP case, most of the loci (except some loci in Ter MD) show lower diffusion mobility compared to the WT. We believe that this mostly results from the fact that the absence of MatP removes the insulation of Ter from the other MDs, which introduces additional contacts between MDs, especially with Ter MD, thereby further slowing down the mobility of chromosomal loci in ΔMatP chromosome. To verify the compactness of Ter MD, we have calculated distance (*D*_*ij*_), between *i*^*th*^ and *j*^*th*^ beads in Ter MD for WT and Δ*MatP* cases, and plotted the distribution of this *D*_*ij*_ in figure S5. From this figure, it is clear that the mean value and standard deviation are large for Δ*MatP* compare to WT, which apprises that Ter MD is less compact for Δ*MatP* compare to WT. We have also reported the total number of connections in our model between different MDs and NS regions in table S1 and table S2 for WT and ΔMatP respectively.

In a bid to investigate how frequent and which particular pair of chromosomal loci meet with each other for the first time. we have calculated first passage time (FPT) for the chromosomal loci from each of the total 200 trajectories. Figure 5 (a) and (b) compare the distribution of FPT for all the loci with all trajectories simulated with Hi-C and without Hi-C interaction respectively. In presence of Hi-C interactions, numerous loci meet each other significantly and more quickly for the first time compared to that in absence of Hi-C, resulting in much shorter mean first passage time (MFPT) of 27.14*τ*_*BD*_. (compared to almost double value in absence of Hi-C). More importantly, a wide distribution of FPT (with long tail) (5 (a)), evident in presence of Hi-C, reflects an inherently heterogenous, Hi-C interaction-modulated encounter between the loci. Figures 5 (c) and (d) demonstrate the network plots of FPT of chromosomal loci in presence and in absence of Hi-C respectively, with the thickness of the lines representing the value of average FPT. The presence of nonspecificity, largely uniform encounters between the loci is clearly evident when the Hi-C interactions are turned off.

**FIG. 5.**
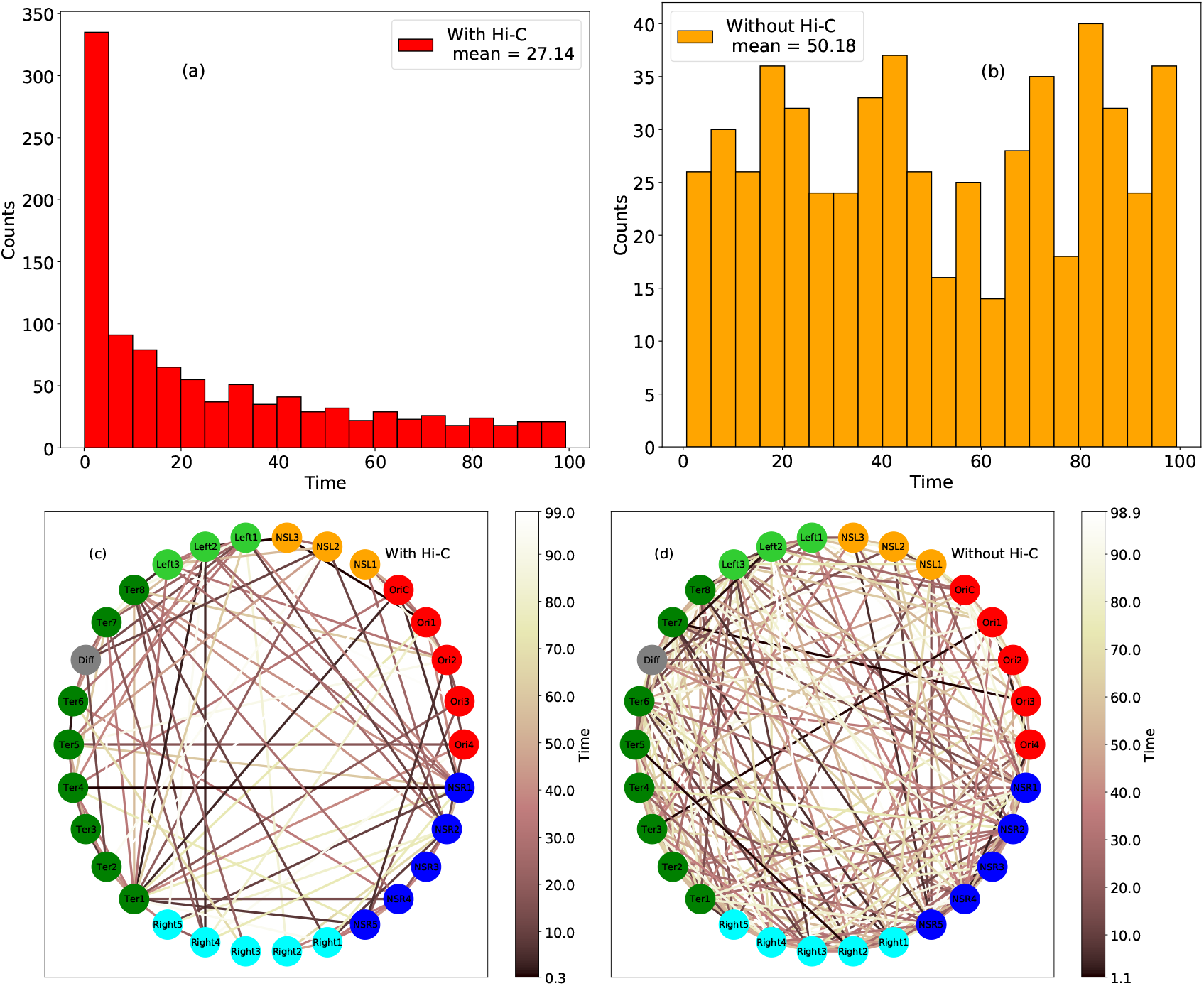
(a), (b) distribution of First passage time (FPT) with all the trajectories in presence of Hi-C, and in absence of Hi-C respectively. For Hi-C cases, the count of first-time meetings at a small time is very large compared to the absence of Hi-C case. (c), (d), represent the network plots of averaged FPT for different loci in presence of HI-C and no Hi-C respectively. Loci are denoted with different colors and placed in a circular orbit according to their genomic co-ordinate sequence and the connection between them represents the value averaged FPT. Color bars are drawn with respect to the value of the averaged FPT. All length and time values are in units of *σ* and *τ*_*BD*_ respectively.

### D. Tracking the origin of sub-diffusive motion: Spatial and Temporal Coherence

Correlation functions are popular approaches, which provide dynamical insights for a system. Mathematically, velocity auto-correlation function (VAF) is defined as, 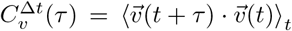, where angular brackets denote the time average and 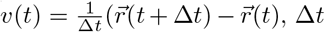, Δ*t* are the different time intervals for velocity calculation and *τ* is the lag time. Physically VAF provides the strength of correlation between two velocities of the particle separated by a time interval *τ*. Here we have calculated an ensemble averaged VAF for the genomic midpoint of the chromosome. Previously experimental and simulations studies on *E. coli* Chromosome [21, 22, 31] have revealed the presence of a negative correlation peak at *τ* = Δ*t* [32], which slowly goes to zero (*τ >>* Δ*T*), due to the viscoelastic nature of cytoplasm. Figure 6 (a) and 6 (b) manifest VAF of *E. coli*. chromosome for small and large time intervals Δ*t* respectively. The observation of negative correlation peak in our model is qualitatively consistent with the previous experiments and implies that our model effectively mimics a viscoelastic cytoplasm, even without explicit incorporation of viscoelasticity as well as a memory kernel [33]. We believe that the presence of long-range inter-bead connectivity in the form of ‘Hi-C’ contacts, is an essential trait of our model, is decisive in furnishing the memory and viscoelasticity in an implicit way. Instead of the genomic midpoint of the chromosome, we have also calculated VAF for Ori2 and Ter3 loci (figure S6 (a), (b), (c), and (d) in SI), which also show the same type of behavior.

**FIG. 6.**
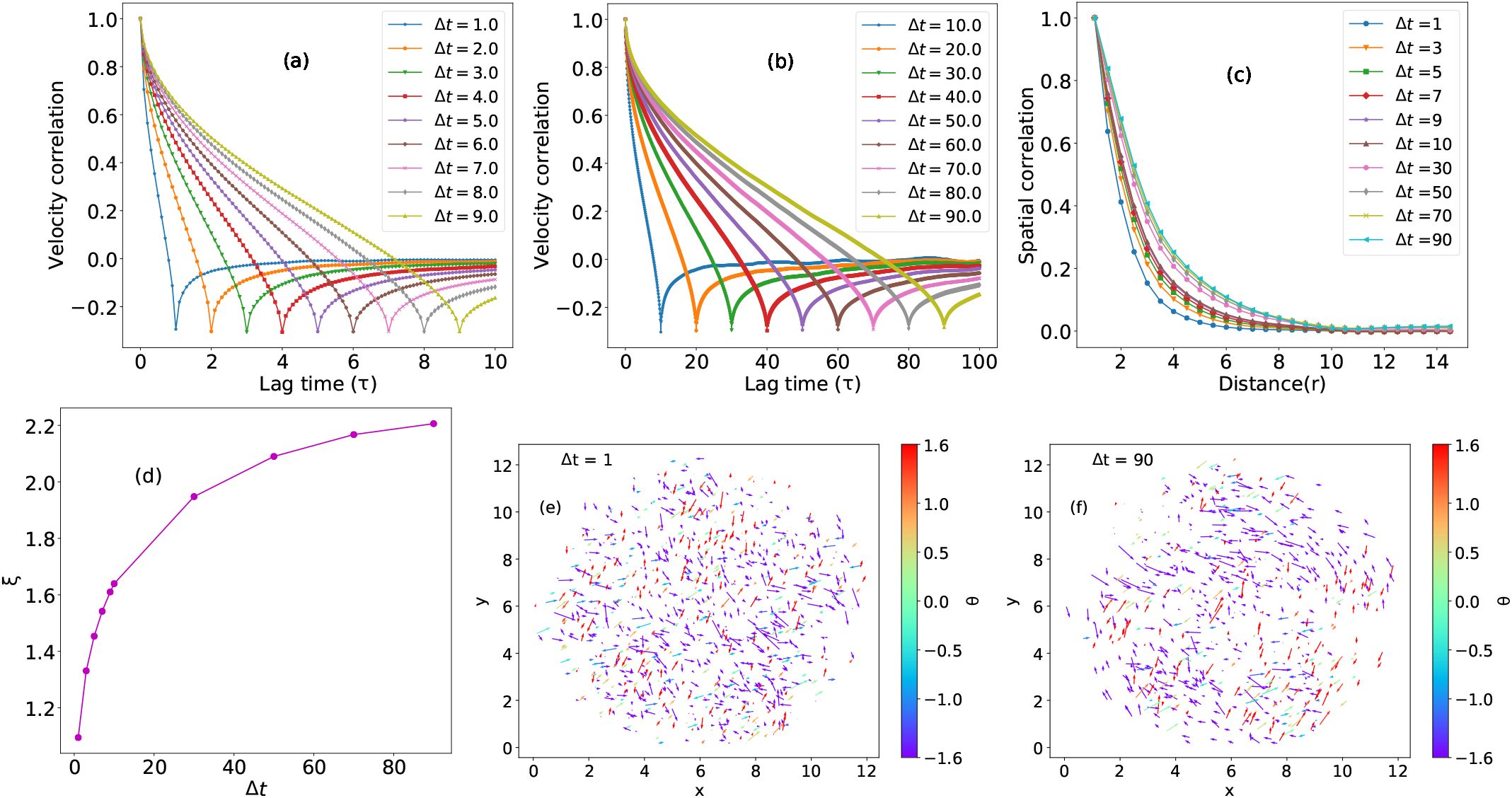
Velocity auto-correlation function(VAF) as a function of lag time for different time interval (a) Δ*t* = (1.0 − 9.0)*τ*_*BD*_ and (b) Δ*t* = (10 − 90)*τ*_*BD*_. Both curve shows that there is a negative correlation peak at *τ* = Δ*t* and slowly goes to zero (*τ >>* Δ*T*), due to the viscoelastic nature of cytoplasm. (c) Spatial correlation 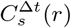 as a function of distance(*r*) for different time interval Δ*t*. As time difference Δ*t* increases, the correlation function decay more slowly. (d) Spatial correlation 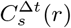 fitted with exponentially decay function as 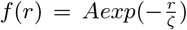 Where A is the constant value and *ζ* is correlation length. *ζ* increases with small values of Δ*t*, but for large values of Δ*t, ζ* values are almost saturated. (e)Vector field of displacement vector for time interval Δ*t* = 1. (f)Vector field of displacement vector for time interval Δ*t* = 90. Color bars represent the angle (*θ*) of the displacement vector. For a small value of time interval, Δ*t* displacement vectors are more irregular compare to large time interval Δ*t*. All length and time values are in units of *σ* and *τ*_*BD*_ respectively.

Motivated by Zidovska et al.’s work [34], where they have used displacement correlation spectroscopy (DCS) to study Spatio-temporal evolution of the global chromatin dynamics in vivo in nuclei of human HeLa cells, we also used the same method for spatial correlation calculation. Spatial correlation of the chromosomal loci defined as (equation 1 [35])

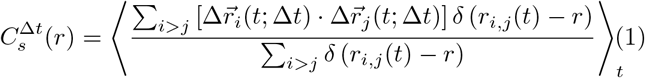

Where *i, j* are the monomer indices, 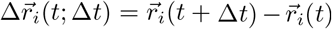 and Δ*t* are the differences in time. Physically this correlation function provides the information on how the displacements of the chromosomal loci, separated by a distance *r*, over time interval Δ*t*, are correlated. Figure 6 (c) shows this spatial correlation 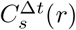 as a function of *r*. As time difference Δ*t* increases, the correlation function decays more slowly. We have also fitted 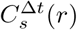 by exponentially decay function as equation 2

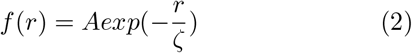

Where A is a constant and *ζ* is the dimension of length, called correlation length (All fitting plots and parameters are given in Figure S7 and table S3 in SI). Figure 6 (d) demonstrates the correlation length (*ζ*) as a function of time difference Δ*t*. For small values of Δ*t, ζ* values are increasing with Δ*t*, but for large values of Δ*t, ζ* values are almost saturated. To quantify the spatial correlation, we have projected the displacement vector 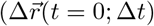 with Δ*t* = 1*τ*_*BD*_ and Δ*t* = 90*τ*_*BD*_) onto the xy plane with a condition (− *σ* ≤ *z* ≤ *σ*), and represented them by a vector field. Figure 6 (e) and 6 (f) depicts the vector field of displacement vector with Δ*t* = 1*τ*_*BD*_ and Δ*t* = 90*τ*_*BD*_ respectively. From these two figures it is clear that displacement vectors are more random for small Δ*t* compared to large Δ*t*, which assist to increase the correlation length with Δ*t*.

### E. The Heterogeneity in loci mobility is robust in presence of active noise

Generally, in normal diffusion, we assumed that the systems are in thermal equilibrium, but in a real scenario, there are numerous ATP-dependent biological activities (transport, metabolism) which prompts the cells to proceed far from equilibrium. A series of numerous past experiments [36–41] on eukaryotic cells and polymer model [42–45], have divulged the role of these biological active processes that initiate non-thermal fluctuations. On a related note, an important experiment by Weber at al. [24] has revealed that ATP-dependent fluctuation (non-thermal) control the *in vivo* dynamics of *E. coli* chromosomal loci. However, the investigation of dynamics of chromosome inside the untreated and ATP-depleted E. coli cells suggested that the sub-diffusive scaling exponent (*α*_*untreated*_ = 0.39, *α*_*treated*_ = 0.40) is almost unchanged irrespective of presence or absence of ATP. However the MSD and *D*_*app*_ (apparent diffusion coefficient) values are significantly reduced in ATP-depleted cell.

In light of present investigations’ observation that chromosomal loci display a heterogeneous and coordinate-dependent mobility, we wanted to explore if this result would be replicated in presence of active noise, which is a better representative of in-vivo environment. To introduce this into our computer model, we increased the strength of the thermal noise and performed a fresh set of dynamical simulations of the chromosome. In particular, the fluctuation dissipation theorem was accordingly changed to 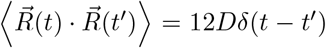 [35, 46–48] instead of the usual expression 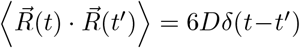. We term our original simulations (which obey fluctuation dissipation theorem) as ‘passive’ and current simulations (which violate fluctuation dissipation theorem) as ‘active’. Figure 7 (a) and figure 7 (b) show the MSD value as a function of lag time for two loci Ori2 and Ter3 respectively, with active and passive noise. From these two figures, it is evident that the MSD values of these loci are notably higher in presence of ‘active’ noise than that of ‘passive’ noise. A more complete comparison across all 30 loci is presented in figure 8. We find that the apparent diffusion constant (*D*_*app*_) of each chromosome locus is significantly increased (almost double) in presence of ‘active noise’ (figure 8 (a-b) in both short((0.1 − 10)*τ*_*Bd*_) and long ((10 − 100)*τ*_*BD*_) time lag respectively. However, very interestingly, as demonstrated by Figure 8 (c) and figure 8 (d), the respective sub-diffusive exponent (*α*) for individual loci remains unchanged in presence of active and passive noise, at both short((0.1 − 10)*τ*_*Bd*_) and long ((10 − 100)*τ*_*BD*_) time lags. Together these results indicate that due to the active noise or ATP-dependent biological activities, mobility of chromosomal loci change remarkably but sub-diffusive exponents are almost fixed. The observation of a robust MSD exponent irrespective of active or passive cytoplasm is consistent with previous experiment and might be due to the fact that viscoelastic nature of cytoplasm, which is effectively captured by our model’s Hi-C interactions, does not change notably even in presence of active noise. Nonetheless, figure 8 also asserts that the inherent heterogeneity and loci-dependence of the mobility and MSD exponents, as found in otherwise ‘passive’ simulations, are rigorously preserved even in presence of ‘active’ noise.

**FIG. 7.**
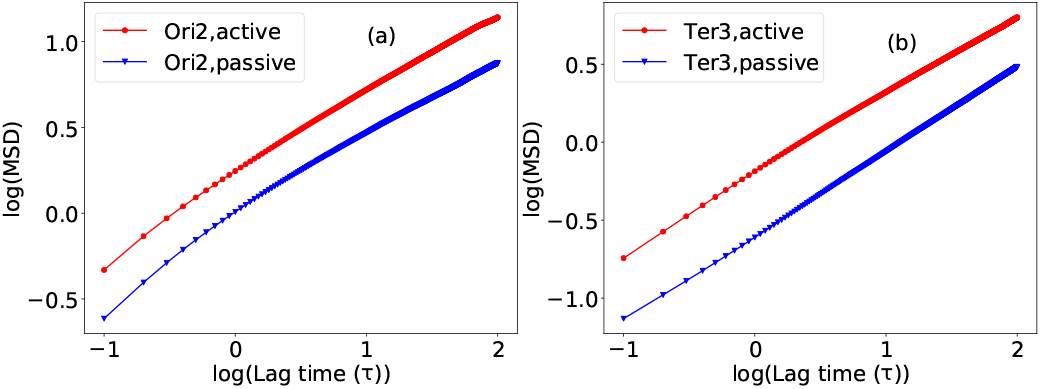
(a) Log-log plot of MSD values as a function of lag time for Ori2 loci with active (red) and passive (blue) noise. (b) Log-log plot of MSD values as a function of lag time for Ter3 loci with active (red) and passive (blue) noise. MSD values of these loci are significantly different when compared with active and passive noise.

**FIG. 8.**
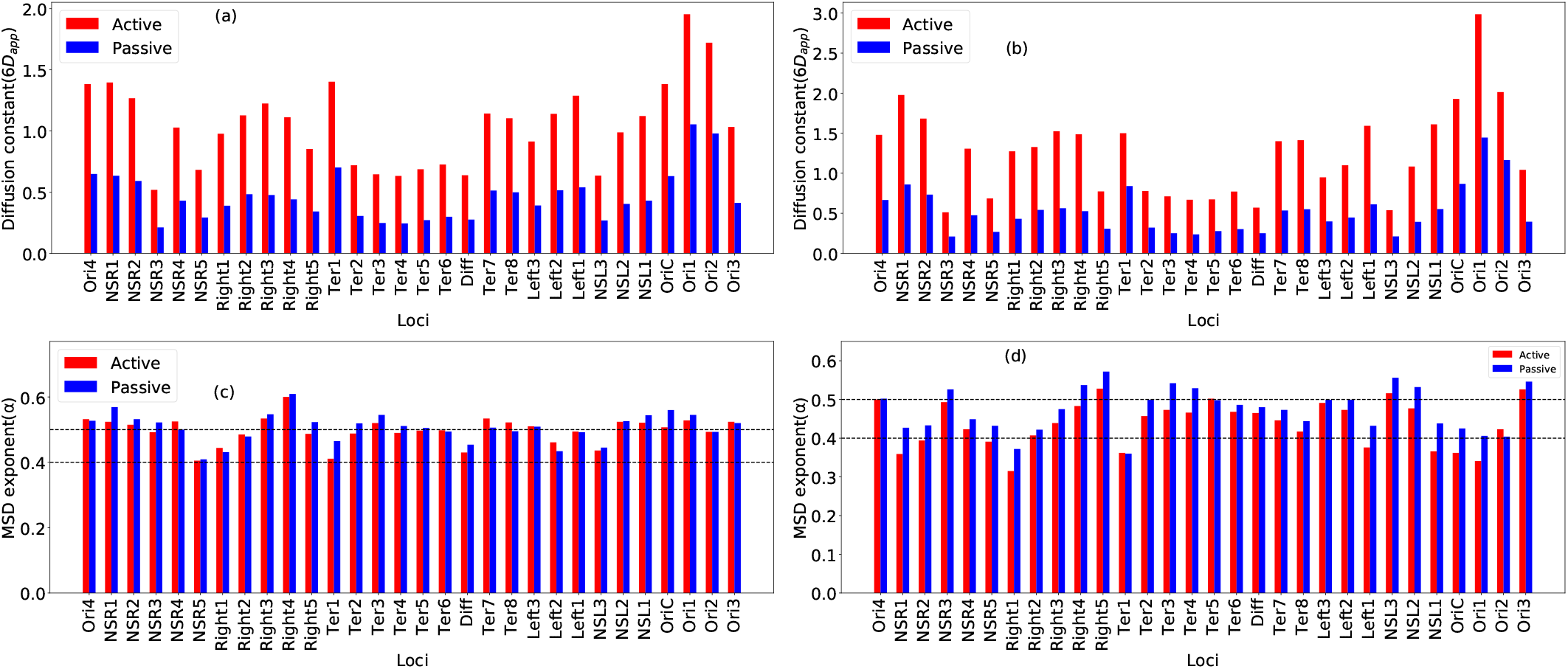
(a) Bar plot of Diffusion constant as a function chromosomal loci for lag time (0.1 − 10)*τ*_*BD*_ in presence of active (red) and passive (blue) noise. (b) Bar plot of Diffusion constant as a function chromosomal loci for lag time (10 − 100)*τ*_*BD*_ in presence of active (red) and passive (blue) noise. For active case Diffusion constant is almost double as compared to the passive case. (c) Bar plot of MSD exponent as a function chromosomal loci for lag time (0.1 − 10)*τ*_*BD*_ in presence of active (red) and passive (blue) noise. (d) Bar plot of MSD exponent as a function chromosomal loci for lag time (10 − 100)*τ*_*BD*_ in presence of active (red) and passive (blue) noise. For both cases (active and passive), sub-diffusive exponents are not changing significantly.

## III. CONCLUDING REMARKS

Our Hi-C data integrated beads-spring polymer model with Brownian dynamics simulations, reveals spatiotemporal dynamics which provides an close-view interpretation of previous experimental and theoretical findings on prokaryotic [21–25] chromosomal loci. We observed that chromosomal loci are moving sub-diffusively but there is clear heterogeneity in sub-diffusive exponents with respect to genomic co-ordinates as well as time. Especially Ori2 and Ter3 differ significantly in their individual displacements. The mobility of each and every loci is crucially dependent on chromosomal co-ordinates. We have also calculated the mean value of sub-diffusive exponent by taking *α* value from each loci and making a histogram. The mean value is *α*_*mean*_ ∼ 0.502 *±* 0.067 for small lag time and *α*_*mean*_ ∼ 0.447 *±* 0.158 for large lag time.

To deduce the origin of the heterogeneous, coordinate-dependent diffusion, we have run a fresh set of control simulations by turning off the Hi-C interaction potentials. We found that, in absence of Hi-C contacts, chromosomal loci are moving homogeneously and their mobility are unchanged with respect to genomic co-ordinates. This observation provides strong evidence of long-range interloci communications, as manifested by the ‘Hi-C interactions’, as one of key modulators of the local dynamics of the bacterial chromosome. The observation of a negative peak in velocity auto correlation function a around particular time difference (*τ* = Δ*t*) brings out the underlying viscoelastic nature effectively furnished by the Hi-C integrated model.

Quite gratifyingly, the sub-diffusive exponents are robust with respect to active and passive noise, while the mobility of loci changes significantly in presence of active noise. This robust MSD exponent indicates that in presence of active noise, viscoelastic nature of cytoplasm, effectively captured by our model interaction, does not change notably.

The self-organization of *E. coli* chromosome into non-overlapping macro-domains has remained a key structural hallmark of this archetypal bacteria. However, how a domain-separated nucleoid would be influencing the dynamics of the chromosome has remained a key question for a while. This is especially complex, considering the intricacies related to segregation and the loci mobilities. In this respect, while the recent experimental demonstration of coordinate-dependent loci movements have been a intriguing discovery, underlying origin has so far been elusive. The current work brings out the importance of inter-loci encounter, statistically represented by Hi-C contacts, in dictating the heterogeneity of the local, coordinate-dependent chromosomal dynamics. The analysis of statistical encounters in the form of first passage time of contacts (figure 5) and the graded extent of macro-domain-dependent fraction of contacts (figure S3), provide a fair idea on how local chromosomal dynamics is governed by the long-range contacts. The dynamical analysis on MatP-devoid chromosome points towards a crucial role that nucleotide-associated proteins (NAPs) might play a role in the local chromosomal dynamics. Overall, the current set of investigations reported in this article enriches our understanding of *E. coli* ‘s chromosomal dynamics, as well as the extent of information Hi-C captures.

Future extension of the present work via introduction of other macro-molecules such as ribosomes and proteins into the model would enable investigations into the diffusion and localization of such macro-molecules [49, 50] at different levels of chromosomal compaction which are brought about by multiple factors such as varying cell sizes [51], induced stress on the cell [49] and mutations.

## IV. MODEL AND METHODS

### A. Model Details

In Hi-C measurement, the cells are in an ensemble of different replication stages with their respective cell cycles. Here we are interested in short-time chromosomal loci dynamics in minimal medium (wt30MM), i.e there is only one single chromosome and no replication fork. We model the *E. coli* chromosome as a bead-spring polymer chain, with each bead representing 5 *×* 10^3^ bp (5 kb), the same as our previous work [14]. This resolution is also the same as Hi-C interaction maps reported by Lioy et al [13]. So the number of beads present in the polymer chain is 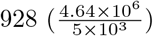.

### B. Interaction Potentials

To mimic the confinement in the *E. coli* cell, we have taken the polymer chain in a spherocyllindrical confinement with average length *L* = 3.08*µm* (including two end caps) and diameter *d* = 0.82*µm*. By taking approximate volume fraction of chromosome *f*_*r*_ = 0.1 [52], we have calculated the bead diameter *σ* to be 0.06731*µm* (SI). Non-bonded interactions of the polymer beads are modeled by the repulsive part of Lenard Jones (LJ) potential i.e *V*_*nb*_(*r*) = 4*ϵ* (*σ/r*)^12^ (Where *ϵ* is in the unit of *k*_*B*_*T* and r is in the unit of *σ*). Bonded interactions between adjacent beads have been modeled by harmonic springs with a spring constant *k*_*spring*_ = 300*k*_*B*_*T/σ*^2^. In a similar fashion, Hi-C contacts are also modeled as harmonic springs with distance dependent force constants and probability dependent bond lengths.

From the Hi-C contact probability matrix we can calculate the distance matrix D as

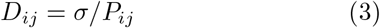

where i and j are row and column index of the matrix respectively. From this distance matrix we define a restraining potential between a pair of Hi-C contacts at a separation of *r*_*ij*_, as 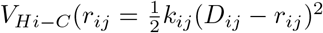. Here *k*_*ij*_ is distance depend force constant that can be calculated from equation 4.

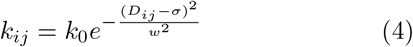

Here P is a sparse matrix with a large number of elements that are 0. So from equation 3, it implies that lot of elements become infinity. To avoid this situation we have modeled *k*_*ij*_ as a Gaussian function. This relation implies that the force constanta becomes smaller for larger distances and the *k*_0_ value is the upper bound of the force constant. We have used a value of *k*_0_ = 10[14]. For optimization of *w* we have calculated the Pearson Correlation coefficient between the experimental and simulated contact probability matrices with varying *w*^2^. Pearson correlation coefficient values show a maximum (88%) at *w*^2^ = 0.3. For the spherocylindrical confinement mimicking the cell wall, we have used a restraining potential in the form of 5.

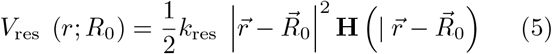

Here *R*_0_ is the center of the spherocylindrical confinement and *k*_*res*_ is the force constant depicting the relative ‘softness’ of the confinement. We have used a value of *k*_*res*_ = 310*k*_*B*_*T/σ*^2^. **H** is a step function that will activate if a polymer bead gets out from the spherocylindrical confinement. Therefore the Hamiltonian can be written as 6

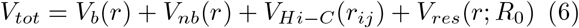

Where *V*_*b*_(*r*), *V*_*nb*_(*r*), *V*_*Hi*−*C*_(*r*_*ij*_), and *V*_*res*_(*r*; *R*_0_) are bonded, non-bonded, Hi-C restraining, and confinement restraining potential respectively.

### C. Simulation Details

To study the dynamics of chromosomal loci, Here we have used Brownian Dynamics (BD) simulations (over-damped condition), by integrating equation 7 (Euler scheme)

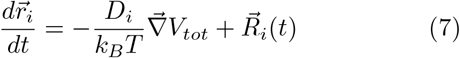

Where *D*_*i*_ is diffusion coefficient of *i*-th bead and 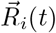 is a random noise, satisfying fluctuation dissipation theorem i.e 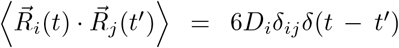. We can calculate *D*_*i*_ from the StokesEinstein equation 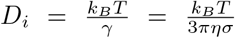, Where *η* is the *E. coli* cytosol viscosity and value is 17.5 Pa-s [53, 54] (26000 times greater than the viscosity of water). Here all simulations time in the unit of 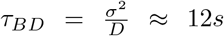 (SI for unit conversion). Integration of the 7 was performed with a time step *δt* = 1 *×* 10^−4^*τ*_*BD*_ by setting *k*_*B*_*T* = 1. Here we have described the simulations details for wt30MM in presence of Hi-C. For wt30MM in absence of Hi-C, ΔMatP and active noise simulation protocol see SI.

All simulations were performed using the open source package GROMACS 5.0.6 [55] and we have modified the source code to introduce the spherocylindrical confinement. We have taken 200 different initial configuration (ensemble) of the polymer chain and minimized the energy of topology by steepest descent algorithm. After energy minimization, we allow the chain for equilibrium with 1 *×* 10^6^*τ*_*BD*_ time steps. Then we collected the data with time step 1 *×* 10^3^*τ*_*BD*_ (data dumping frequency).

## Supporting information

Supplemental informations

## V. ACKNOWLEDGMENTS

We sincerely acknowledge Dr. V. Lioy for providing us the average cell size of the cells grown in MM at 30°C used for their study. We acknowledge the computation facilities provided by TIFR Centre for Interdisciplinary Sciences, India. We acknowledge the support of the Department of Atomic Energy, Government of India, under Project Identification No. RTI 4007.

