## Supplemental informations for "Hi-C Contacts Encode Heterogeneity in Sub-diffusive Motion of *E. coli* Chromosomal Loci"

### 0.1 Calculation of Bead Diameter

For *E. Coli* chromosome at $30^{\circ}C$ in minimal medium (wt30MM), cell length($L$) (including two caps) and diameter($d$) become $3.08\mu m$ and $0.82\mu m$ respectively. Let the radius of end cap be $r = d/2$, and the length excluding end cap be $l = L - d$, diameter of each bead be $\sigma$ and total number of beads be $N = 928$. Now volume of the whole chromosome is $V_{chro} = \frac{4}{3}\pi\sigma^3 N$ and volume of the cell is $V_{cell} = \pi r^2 l + \frac{4}{3}\pi r^3$. Now volume fraction ($f_r$) defined as

$$f_r = \frac{V_{chor}}{V_{cell}} = \frac{\frac{4}{3}\pi\sigma^3 N}{\pi r^2 l + \frac{4}{3}\pi r^3} \tag{1}$$

After rearranging 1 we can calculate $sigma$ as

$$\sigma = \Big(\frac{6 f_r r^2}{N}(\frac{4}{3}r + l)\Big)^{\frac{1}{3}} \tag{2}$$

By taking $f_r = 0.1$ [1] and all other parameter as given above, we have gotten $\sigma = 0.06731\mu m$.
By taking $\sigma = 1$, the length and diameter become, $L = 45.754\sigma$ and $d = 12.181\sigma$ respectively in reduced unit (in terms of $\sigma$). In our simulation data all length are in units of $\sigma$

### 0.2 Model and Methods

In the main text, we have described the simulation details for wt30MM in presence of Hi-C. Here we will describe others three cases.

#### 0.2.1 wt30MM(without Hi-C):–

Simulation protocol is same as wt30MM(with Hi-C), but only different is that now $V_{tot}$ has been changed by removing $V_{Hi-C}$ term. The form of $V_{tot}$ is given by equation 3

$$V_{tot}^{noHi-C} = V_b(r) + V_{nb}(r) + V_{res}(r; R_0) \tag{3}$$

Where $V_b(r)$, $V_{nb}(r)$, and $V_{res}(r; R_0)$ are bonded, non-bonded, and confinement restraining potential respectively.

#### 0.2.2 ΔMatP:–

Simulation protocol is same as wt30MM(with Hi-C), but only different is that now $V_{tot}$ has been changed. The form of $V_{tot}$ is given by equation 4

$$V_{tot}^{\Delta MatP} = V_b(r) + V_{nb}(r) + V_{Hi-C}^{\Delta MatP}(r_{ij}) + V_{res}(r; R_0) \tag{4}$$

Where $V_b(r)$, $V_{nb}(r)$, $V_{Hi-C}^{\Delta MatP}(r_{ij})$, and $V_{res}(r; R_0)$ are bonded, non-bonded, Hi-C restraining, and confinement restraining potential respectively. Here $V_{Hi-C}^{\Delta MatP}(r_{ij})$ has been calculated in the same way as done for wt30MM, by taking Hi-C data of mutated bacteria ΔMatP [2].

#### 0.2.3 wt30MM (Active):–

Simulation protocol is same as wt30MM(with Hi-C), but only different is that here random noise does not follow fluctuation-dissipation theorem. The fluctuation dissipation theorem has been changed to $\left\langle \vec{R}(t) \cdot \vec{R}(t') \right\rangle = 12D\delta(t-t')$ instead of the usual expression $\left\langle \vec{R}(t) \cdot \vec{R}(t') \right\rangle = 6D\delta(t-t')$.

### 0.3 Data Analysis

#### 0.3.1 MSD calculation:–

In our simulation, we have taken 200 different initial configurations of the polymer chain (ensemble). The time-average MSD for a particular loci (n) calculated from a single trajectory as $MSD_n(\tau) = \langle \mid \vec{r}_n(t+\tau) - \vec{r}_n(t) \mid^2 \rangle_t = \frac{1}{T_{tot}} \sum_t \mid \vec{r}_n(t+\tau) - \vec{r}_n(t) \mid^2$, where $\tau$ and $T_{tot}$ are lag time and total simulation time respectively. Ensemble averaged MSD for particular loci $n$ has been calculated by averaging over all 200 different configurations and denoted as $(\overline{MSD_n})$. In general MSD follows the equation, $(\overline{MSD_n}) = 6D_{app}\tau^{\alpha}$ (in $3d$), $D_{app}$ is the apparent diffusion coefficient, and $\alpha$ is the MSD exponent. We have fitted our ensemble averaged MSD with this equation and calculated this $6D_{app}$ and $\alpha$ values for two different time lag $(0.1-10)\tau_{BD}$ and $(10-100)\tau_{BD}$.

#### 0.3.2 Velocity auto-correlation:–

We have calculated velocity auto-correlation function by using the equation, $C_v^{\Delta t}(\tau) = \langle \vec{v}(t+\tau) \cdot \vec{v}(t) \rangle_t$, where angular bracket denotes the time average and $v(t) = \frac{1}{\Delta t}(\vec{r}(t+\Delta t) - \vec{r}(t)$, $\Delta t$ is the different time interval for velocity calculation and $\tau$ is the lag time. Here we have also done a ensemble average over all 200 trajectories.

#### 0.3.3 Spatial correlation:–

We have calculated spatial correlation function by using the equation 5,

$$C_s^{\Delta t}(r) = \left\langle \frac{\sum_{i>j} \left[\Delta \vec{r}_i(t;\Delta t) \cdot \Delta \vec{r}_j(t;\Delta t)\right] \delta\left(r_{i,j}(t) - r\right)}{\sum_{i>j} \delta\left(r_{i,j}(t) - r\right)} \right\rangle_t \tag{5}$$

Where $i$, $j$ are the monomer index, $\Delta\vec{r}_i(t;\Delta t) = \vec{r}_i(t+\Delta t) - \vec{r}_i(t)$ and $\Delta t$ is the difference in time. Similarly, here we have also done a ensemble average over all 200 trajectories.

#### 0.3.4 First Passage Time (FPT):–

We have calculated first passage time (FPT) for the chromosomal loci from each of the total 200 trajectories. In our simulation diameter of each bead is $\sigma$. So for FPT calculation, we have calculated the distance between all the loci for each time interval data and we have also used a distance cut-off ($r < 1.1\sigma$). Once the distance between two loci falls within the cut-off, then the corresponding times are recorded. To get better insight into which chromosomal loci meet with each other for the first time and what is their average time for the meeting, we have used a network plot to represent it. First, we have recorded the time and loci index once they meet with each other as done previously.

If the same loci meet in different trajectories then we have calculated a averaged first passage time by averaging over FPT from different trajectories for a repeated index of loci.

### 0.4 Unit Conversion

For simulation purpose we have used $\epsilon$ in the unit of $k_BT$ and time in the unit of $\tau_{BD}$. From Einstein-Stokes relation, diffusion co-efficient of a bead is $D = \frac{k_BT}{\gamma} = \frac{k_BT}{3\pi\eta\sigma}$, where $K_B$ is Boltzmann constant, T is temperature, $\eta$ is viscocity of the medium and $\sigma$ is diameter f bead. Using this D value we can calculate Brownian time scale $\tau_{BD}$ as $\tau_{BD} = \frac{\sigma^2}{D} = \frac{3\pi\eta\sigma^3}{k_BT} \simeq 12s$ (in real units, by taking real value $T = 303$K, $\eta = 17.5$ Pa-s[3, 4] and $\sigma = 0.06731\mu m$).

### 0.5 Distribution of MSD for loci Ori2 and Ter3

We have calculated the MSD value from each trajectory for Ori2 and Ter3 loci, at a particular lag time ($15\tau_{BD}$) and made a distribution of this MSD value. Figure S1 (a) and figure S1 (b) shows the distribution of MSD value for Ori2 and Ter3 loci respectively. MSD value for Ori2 ($(1.123 - 5.541)\sigma^2$) are very large compared to Ter3 ($(0.567 - 1.720)\sigma^2$). This difference value in MSD clearly suggest that loci mobility depend on chromosomal coordinate.

### 0.6 Apparent diffusion coefficient for all loci in absence and in presence of Hi-C

We have calculated apparent diffusion constant by fitting the MSD values as $(\overline{MSD_n}) = D\tau^\alpha$ and denoted as $D = 6D_{app}$. Figure S2 (a) and Figure S2 (b) shows the apparent diffusion coefficient value as a function of chromosomal coordinate, in presence of Hi-C with lag time ($(0.1 - 10)\tau_{BD}$) and ($(10 - 100)\tau_{BD}$) respectively. Similarly Figure S2 (c) and Figure S2 (d) depicts the apparent diffusion coefficient value as a function of chromosomal coordinate, in absence of Hi-C with lag time ($(0.1 - 10)\tau_{BD}$) and ($(10 - 100)\tau_{BD}$) respectively. In absence of Hi-C all loci shows nearly uniform value of diffusion constants. In presence of Hi-C there is a clear heterogeneity in apparent diffusion constants.

### 0.7 Fraction of Contacts

We have calculated the number of connections between different beads and normalized them between $(0 - 1)$ and plotted a heat map. Figure S3 shows the heat map of the fraction of Hi-C contact formed within each of the MDs and NS regions. The fraction of connections in Ter MD is high compare to other MDs and NS regions, which makes the Ter MD more compact.

### 0.8 Comparison of diffusion constant between WT and $\Delta$MatP

Figure S4 (a) and (b) depicts the variation of diffusion constant as a function of chromosomal loci for two lag time ($(0.1 - 10)\tau_{BD}$) and ($(10 - 100)\tau_{BD}$) respectively. In $\Delta$MatP case most of the loci except some of the loci in Ter MD, show less diffusion constants as compared to the WT case.

### 0.9 Distribution of Distance $D_{ij}$

We have calculated the distance $D_{ij}$ between $i^{th}$ and $j^{th}$ beads in Ter MD and plotted a distribution of this $D_{ij}$ values. Figure S5 shows the mean value and standard deviation is large for $\Delta$MatP compare to WT, Which tells that Ter MD is less compact in absence of MatP. We have also reported the total number of connection in table S1 and table S2. Although the number of connections for $\Delta$MatP in Ter MD is large compare to WT, but effective distances $D_{ij}$ (figure S5) between $i^{th}$ and $j^{th}$ beads increases for $\Delta$MatP, which provides less close-packed.

### 0.10 Velocity Auto-correlation Function (VAF) for Ori2 and Ter3 loci

Here we have calculated velocity auto correlation function (VAF) for Ori2 and Ter3 loci in a similar way as we calculated for the genomic midpoint. Both Ori2 and Ter3 loci also show similar kind of viscoelastic property i.e there is a negative peak at $\tau = \Delta t$ and slowly goes to zero ($\tau >> \Delta T$). Figure S6(a) and figure S6(b) shows the (VAF) for Ori2 with time difference $\Delta t = (1.0 - 9.0)\tau_{BD}$ and $\Delta t = (10 - 90)\tau_{BD}$ respectively. Similarly figure S6(c) and figure S6(d) depicts the (VAF) for Ter3 with time difference $\Delta t = (10 - 90)\tau_{BD}$ and $\Delta t = (10 - 90)\tau_{BD}$ respectively.

### 0.11 Fitting Parameter for Correlation Length

Figure S7 demonstrate fitting of spatial correlation function with exponential decay function ($f(r) = Aexp(-\frac{r}{\zeta})$). Table S3 provides all fitting parameter. Correlation length increases with time interval but almost saturated for larger time intervals.

### 0.12 Essential software and libraries

For the simulations, we have used GROMACS 5.0.6 [5] (a molecular dynamics simulation package) and for data analysis, we have used our own homemade Python [6] scripts. For programming, plotting, and fitting, we have used Matplotlib [7], NumPy [8], SciPy [9], and NetworkX [10]. For visualization of the simulated trajectories and for rendering , we have used Visual Molecular Dynamics (VMD) [11] Software. For other scientific illustrations, we have used Inkscape [12].

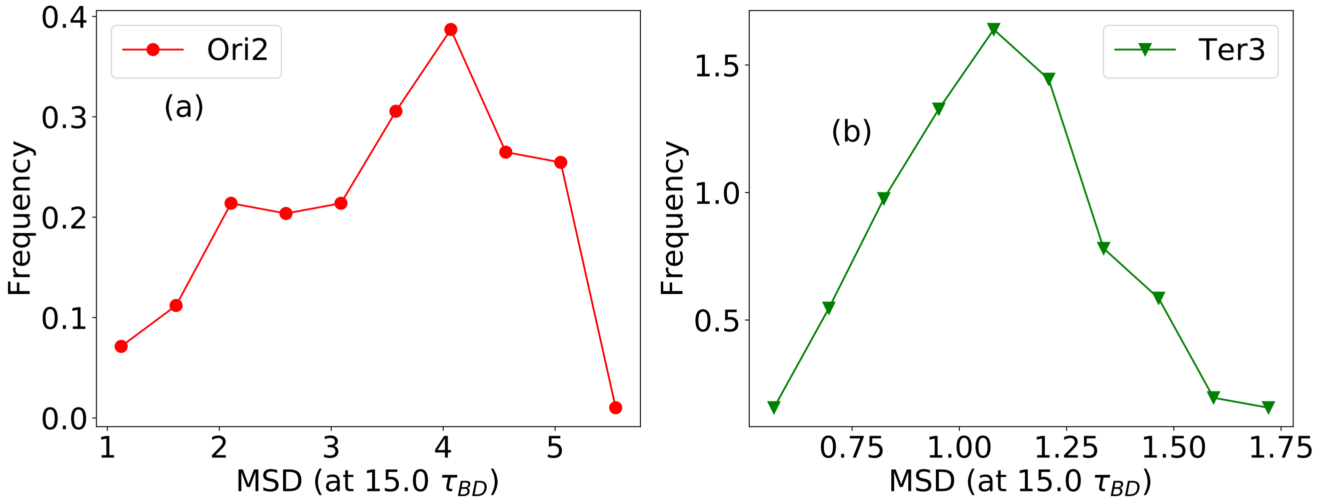


Figure S1: (a) Distribution of MSD values for Ori2 loci with lag time $15\tau_{BD}$ (b) Distribution of MSD values for Ter3 loci with lag time $15\tau_{BD}$. MSD values for Ori2 ($(1.123 - 5.541)\sigma^2$) are very large compare to Ter3 ($(0.567 - 1.720)\sigma^2$).

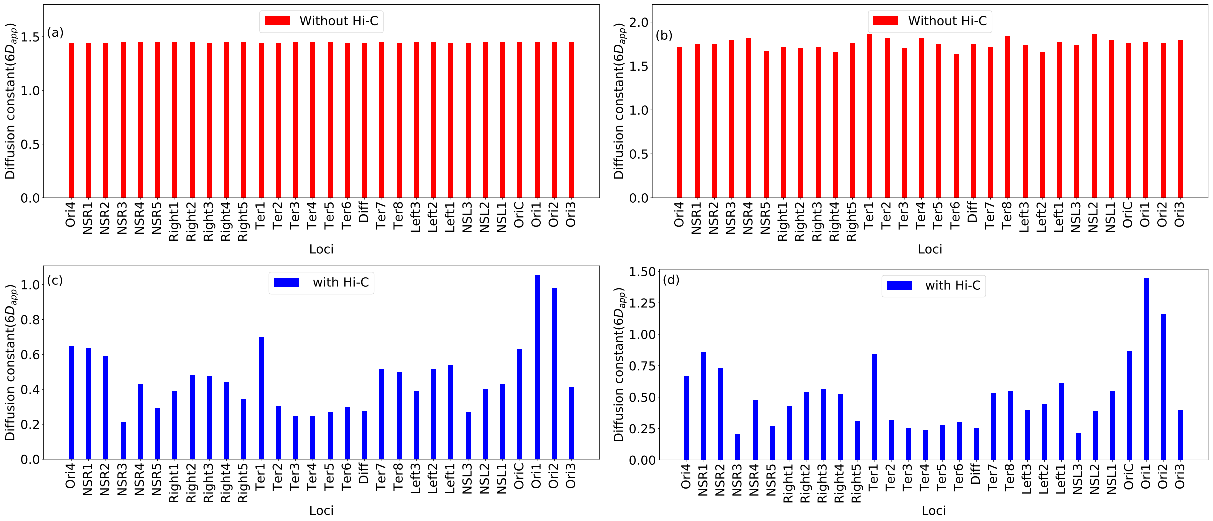


Figure S2: Bar plot of diffusion constant as a function chromosomal loci in absence of Hi-C with lag time (a) $(0.1-10)\tau_{BD}$ and (b) $(10-100)\tau_{BD}$. In both case the values of diffusion constant are same, so mobility are same for all the loci. Bar plot of Diffusion constant as a function chromosomal loci in presence of Hi-C with (c) lag time $(0.1-10)\tau_{BD}$ and (d) $(10-100)\tau_{BD}$. In both case the values of diffusion constant vary from loci to loci, so mobility of chromosomal loci depend on their gnomic co-ordinate. For all the plot Diffusion constants are unit of $\frac{\sigma^2}{\tau_{BD}^{\alpha}}$.

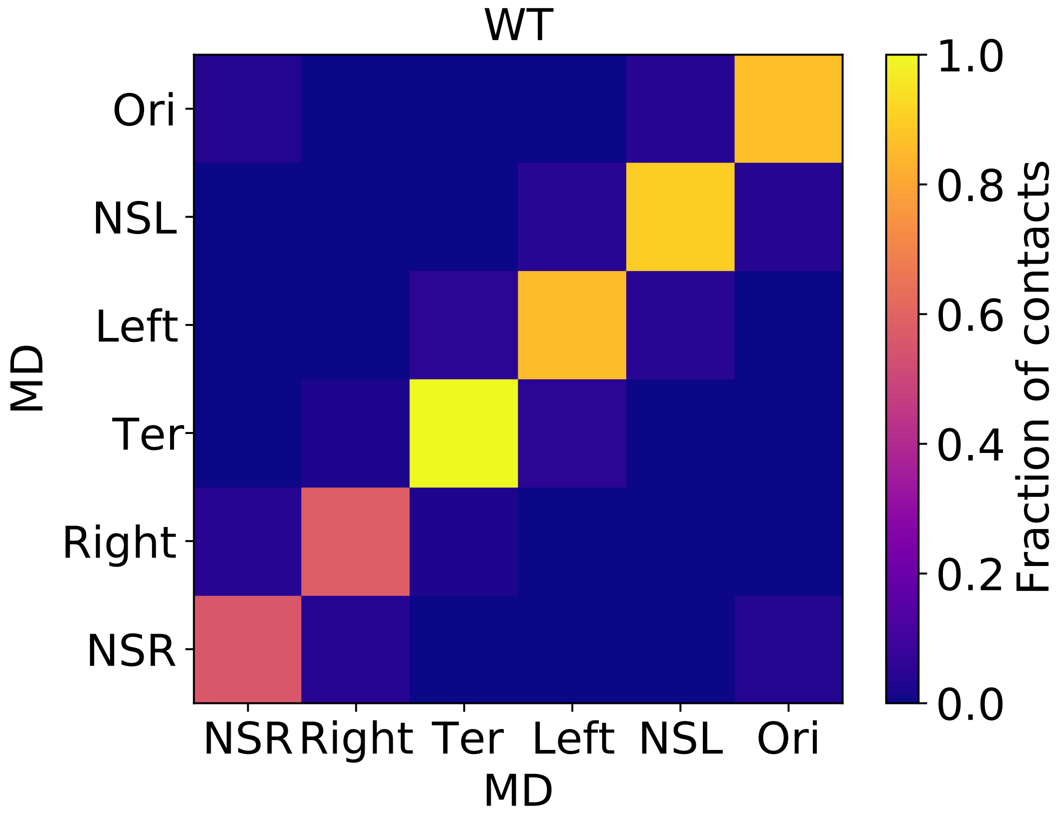


Figure S3: Heat map of fraction of contacts between inter and intrachromosomal loci for different MDs and NS regions. Ter MD shows more intrachromosomal interaction compare to others MDs and NS regions.

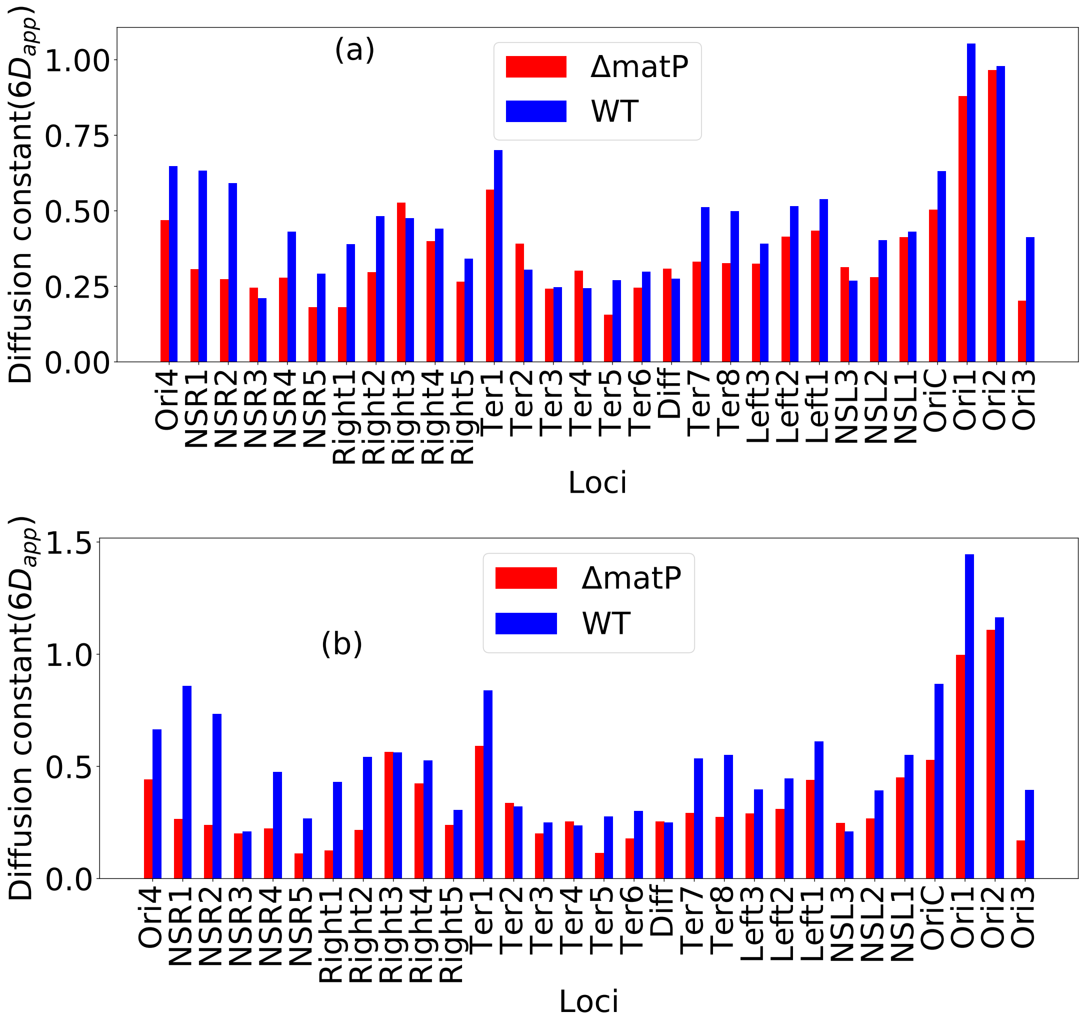


Figure S4: Comparison of diffusion constant values with wild type (WT) and mutant (ΔmatP) cases for two different lag time (a) $(0-10)\tau_{BD}$ and (b) $(10-100)\tau_{BD}$. For mutant case, most of the loci show less diffusion constant value compare to WT case.

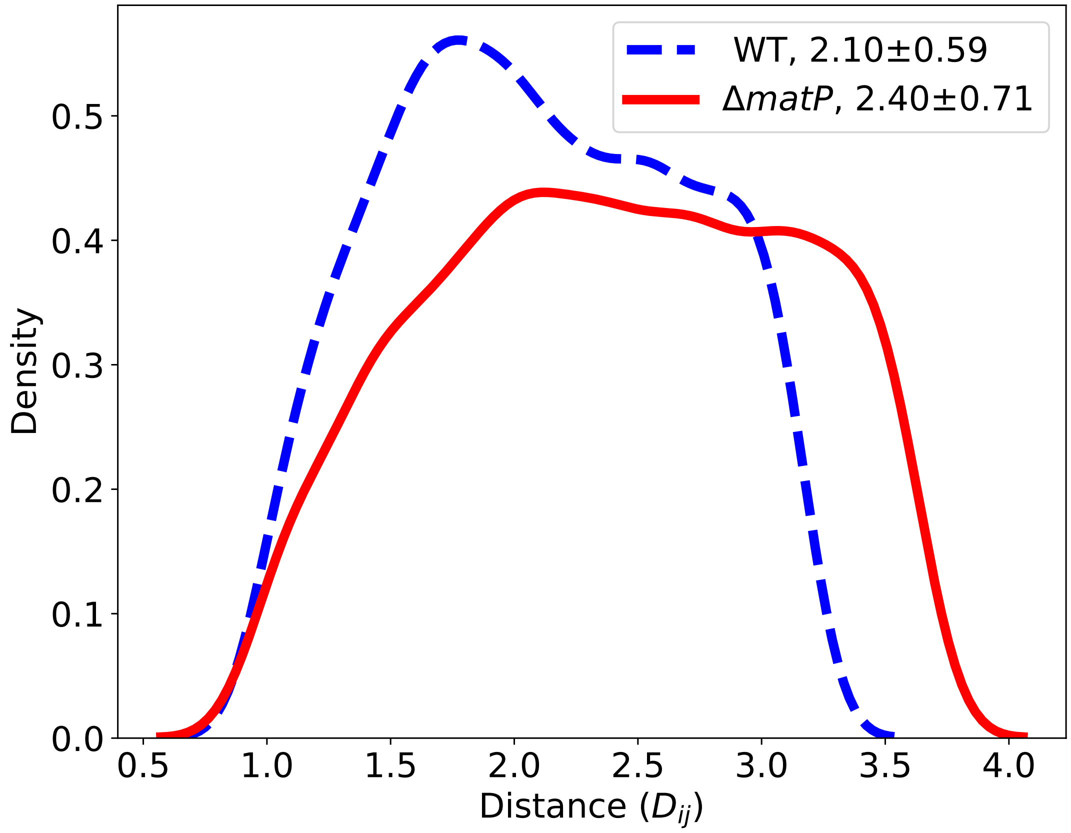


Figure S5: Distribution of distance ($D_{ij}$) between two beads in Ter MD, in presence and absence of MatP. For ΔMatP case mean value and standard deviation is large compare to WT case, which confirms the less compactness of Ter region for ΔMatP case compare to WT cases.

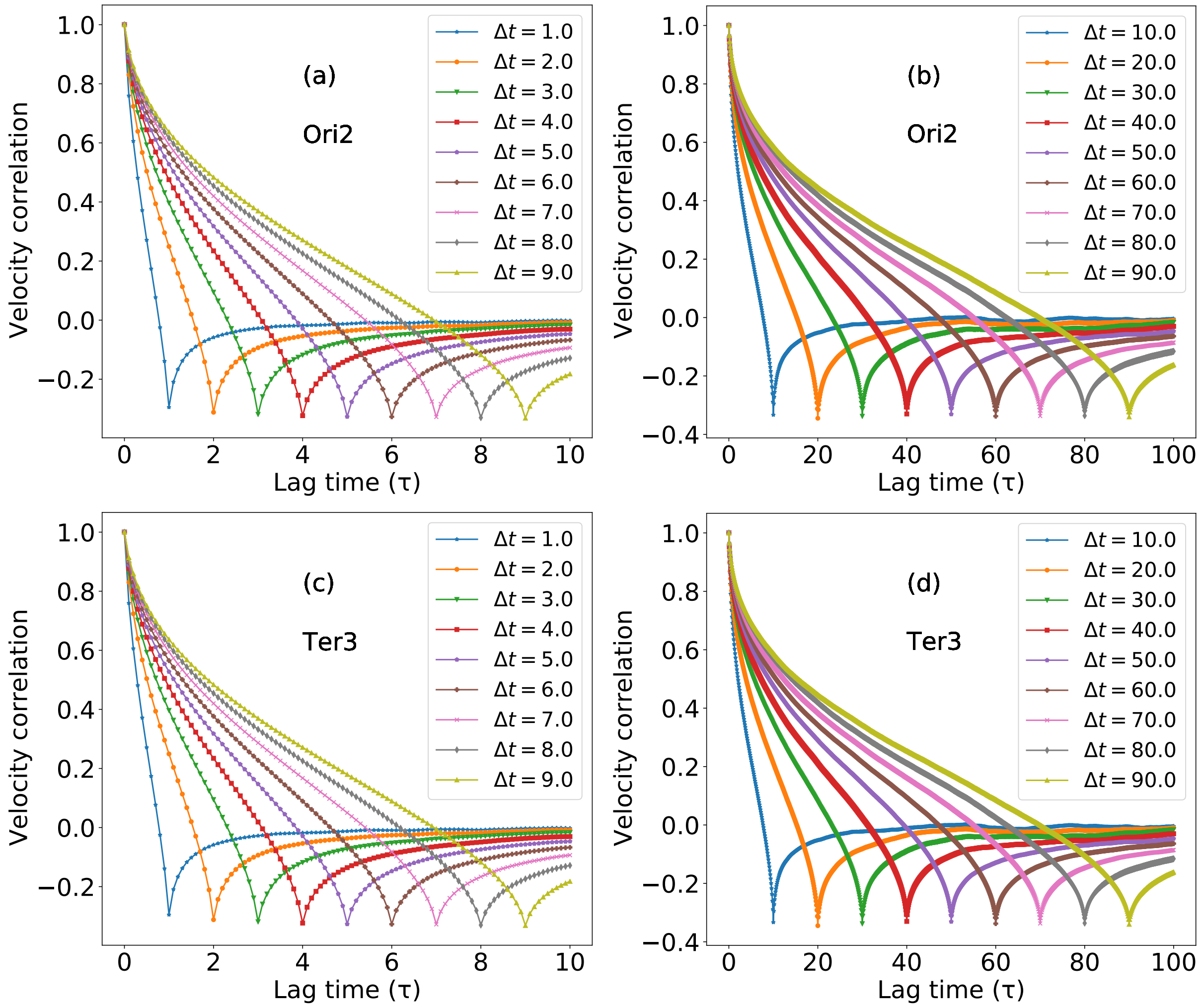


Figure S6: (a) velocity auto-correlation function(VAF) as a function of lag time with different time interval $\Delta t = (1.0 - 9.0)\tau_{BD}$ for Ori2 loci. (b) velocity auto-correlation function(VAF) as a function of lag time with different time interval $\Delta t = (10 - 90)\tau_{BD}$ for Ori2 loci. (c) velocity auto-correlation function(VAF) as a function of lag time with different time interval $\Delta t = (1.0 - 9.0)\tau_{BD}$ for Ter3 loci. (d) velocity auto-correlation function(VAF) as a function of lag time with different time interval $\Delta t = (10 - 90)\tau_{BD}$ for Ter3 loci.

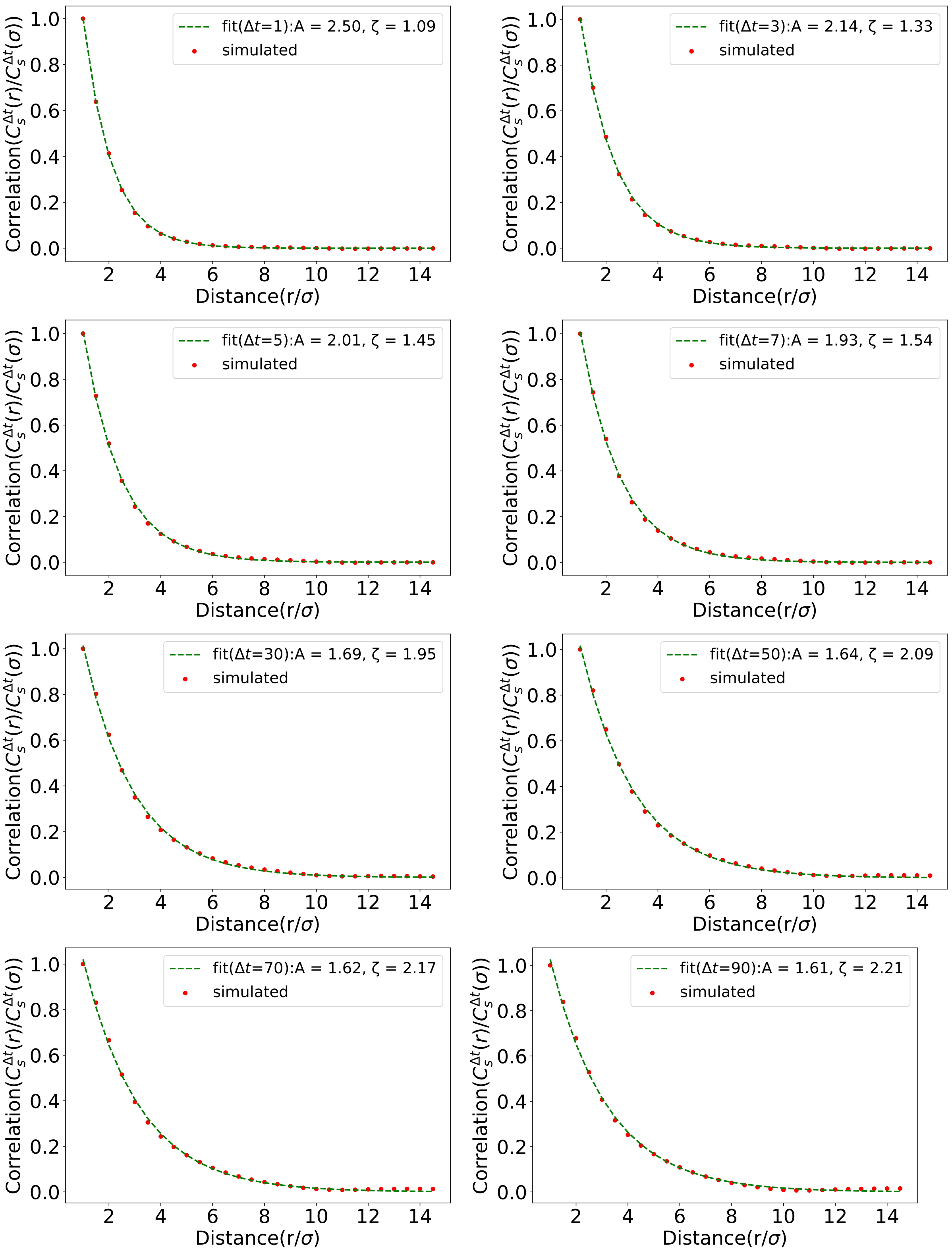


Figure S7: Spatial correlation are fitted with exponential decay function for different time interval $\Delta t$. Correlation length increases with time interval but almost saturated for larger time interval.

| WT | NSR | Right | Ter | Left | NSL | Ori |
|---|---|---|---|---|---|---|
| NSR | 3680 | 297 | 1 | 1 | 1 | 241 |
| Right | 297 | 3828 | 172 | 0 | 0 | 0 |
| Ter | 1 | 172 | 6574 | 341 | 0 | 0 |
| Left | 1 | 0 | 341 | 5634 | 318 | 0 |
| NSL | 1 | 0 | 0 | 318 | 5916 | 275 |
| Ori | 241 | 0 | 0 | 0 | 275 | 5676 |

Table S1: No of connections between different MDs and NS regions for WT.

| $\Delta$MatP | NSR | Right | Ter | Left | NSL | Ori |
|---|---|---|---|---|---|---|
| NSR | 4896 | 483 | 1 | 0 | 1 | 380 |
| Right | 483 | 4206 | 240 | 0 | 1 | 1 |
| Ter | 1 | 240 | 7462 | 447 | 0 | 0 |
| Left | 0 | 0 | 447 | 6774 | 419 | 0 |
| NSL | 1 | 1 | 0 | 419 | 6982 | 343 |
| Ori | 380 | 1 | 0 | 0 | 343 | 6758 |

Table S2: No of connections between different MDs and NS regions for $\Delta$MatP.

| Time interval ($\Delta t$) | $A$ | $\zeta$ |
|---|---|---|
| 1.0 | 2.50 | 1.09 |
| 3.0 | 2.14 | 1.33 |
| 5.0 | 2.01 | 1.45 |
| 7.0 | 1.93 | 1.54 |
| 9.0 | 1.88 | 1.61 |
| 10.0 | 1.86 | 1.64 |
| 30.0 | 1.69 | 1.95 |
| 50.0 | 1.64 | 2.09 |
| 70.0 | 1.62 | 2.17 |
| 90.0 | 1.61 | 2.21 |

Table S3: All fitting parameter for exponential decay fitting.

[10] Aric Hagberg, Pieter Swart, and Daniel S Chult. Exploring network structure, dynamics, and function using networkx. Technical report, Los Alamos National Lab.(LANL), Los Alamos, NM (United States), 2008.

[11] William Humphrey, Andrew Dalke, and Klaus Schulten. Vmd: visual molecular dynamics. *Journal of molecular graphics*, 14(1):33–38, 1996.

[12] Inkscape Project. Inkscape.
